# Fast retrograde access to projection neuron circuits underlying vocal learning in songbirds

**DOI:** 10.1101/2020.02.19.955773

**Authors:** DN Düring, F Dittrich, MD Rocha, RO Tachibana, C Mori, K Okanoya, R Boehringer, B Ehret, BF Grewe, M Rauch, JC Paterna, R Kasper, M Gahr, RHR Hahnloser

**Affiliations:** Institute of Neuroinformatics, University of Zürich/ETH Zürich, Zurich, Switzerland; Neuroscience Center Zurich (ZNZ), Zurich, Switzerland; Department of Behavioural Neurobiology, Max Planck Institute for Ornithology, Seewiesen, Germany; Department of Life Sciences, The University of Tokyo, Tokyo, Japan; Viral Vector Facility (VVF) of the Neuroscience Center Zurich (ZNZ), Zurich, Switzerland; Imaging Center of the Max Planck Institute of Neurobiology, Munich, Germany

## Abstract

Understanding the structure and function of neural circuits underlying speech and language is a vital step towards better treatments for diseases of these systems. Songbirds, among the few animal orders that share with humans the ability to learn vocalizations from a conspecific, have provided many insights into the neural mechanisms of vocal development. However, research into vocal learning circuits has been hindered by a lack of tools for rapid genetic targeting of specific neuron populations to meet the quick pace of developmental learning. Here, we present a new viral tool that enables fast and efficient retrograde access to projection neuron populations. In zebra finches, Bengalese finches, canaries, and mice, we demonstrate fast retrograde labeling of cortical or dopaminergic neurons. We further demonstrate the suitability of our construct for detailed morphological analysis, for *in vivo* imaging of calcium activity, and for multicolor brainbow labeling.

## Introduction

Speech and language disorders affect millions of children worldwide^1^. These disorders are associated with a number of pathologies, including autism spectrum^2^ and attention deficit hyperactivity disorders^3^, but they also occur in otherwise normally developing children. The causes of speech and language disorders are poorly understood, but abnormal brain structure likely plays a crucial role^4^□. Interestingly, neither mice, the most common animal model for pre-clinical studies, nor our closest relatives, non-human primates, share the human ability to learn vocalizations by imitating a conspecific^5^. Songbirds, on the other hand, are gifted vocal learners and display many parallels with speech acquisition in humans^6^□. Therefore, songbirds provide an excellent model system to examine the neural circuits associated with speech development. Songbird circuit research remains, however, limited by the current lack of transgenic animal lines and proper neurogenetic tools.

The virally mediated delivery of genetic cargo to a specific neuron population is a highly valuable approach for probing anatomy and circuit function. By entering at axon terminals, retrograde viral vectors can specifically transduce projection neuron populations and provide direct access to the wiring and function of the brain^7,8^. Unfortunately, viral vectors are not easily transferable between model species. For example, rabies and canine adenovirus (CAV), the most commonly used retrograde vectors in rodents, have never been shown to successfully infect avian tissue. Other examples include adeno associated viral (AAV) vectors that are commonly used in rodents and other mammals, but suffer from major limitations in accessing songbird projection neurons^9–15^.

The limitations of the AAV vectors so far applied for retrograde access to songbird projection neurons are four-fold. First, these AAV vectors result in low transduction success rates, with only 18% to 43% of intracerebral injections resulting in successful transgene expression^9^. Second, their transduction of projection neurons is sparse, resulting in transgene expression in small portions of the targeted populations^9,11^. This limitation makes these tools incompatible with studies requiring genetic manipulations of a majority of neurons among a given type. The third limitation is weak transgene expression^9,13^. For example, when used to induce fluorescent protein expression, the resulting limited dendritic and axonal labeling^9,13^ makes current retrograde tools unsuitable for morphological studies. Finally, the fourth limitation of the current AAV vectors is that they require long incubation periods of at least 4 to 6 weeks (and up to 6 months) post-injection until transgene expression becomes detectable^10,11,13^. Such long incubation periods typically hinder any vocal development studies in zebra finches (*Taeniopygia guttata*), the most common songbird model species in neuroscience. This is because zebra finches undergo short song learning phases during the critical period for vocal learning. This period ends 90 days after hatching and is accompanied by fast brain rewiring^16^. The rapid succession of vocal learning events in songbirds thus requires fast viral tools for circuit investigations. To date, no viral vector has proven to efficiently target projection neurons of the songbird brain resulting in robust transgene expression in a relevant time frame.

Here we present a fast and reliable retrograde viral tool for the songbird brain. Briefly, we combined a self-complementary (sc) plasmid organization of a capsid variant of AAV-DJ, AAV-DJ/9^17^, and an expression cassette under the transcriptional control of the human cytomegalovirus (hCMV) promoter/immediate-early enhancer (IE) in combination with a chimeric intron (chI), resulting in AAV vectors according to scAAV-2-DJ/9-hCMV-I/E-chI-transgene-p(A) (Fig. 1a,b; Methods). In several songbird species and in mice, we found that our viral construct is efficient in retrogradely transducing projection neuron circuits within about one week after vector injection (Fig. 1c-o; Fig. 2). Furthermore, our construct is suitable for detailed morphological investigations, for multi-color brainbow imaging, and for *in-vivo* recordings of neural activity using fluorescent calcium sensors.

**Fig. 1.**
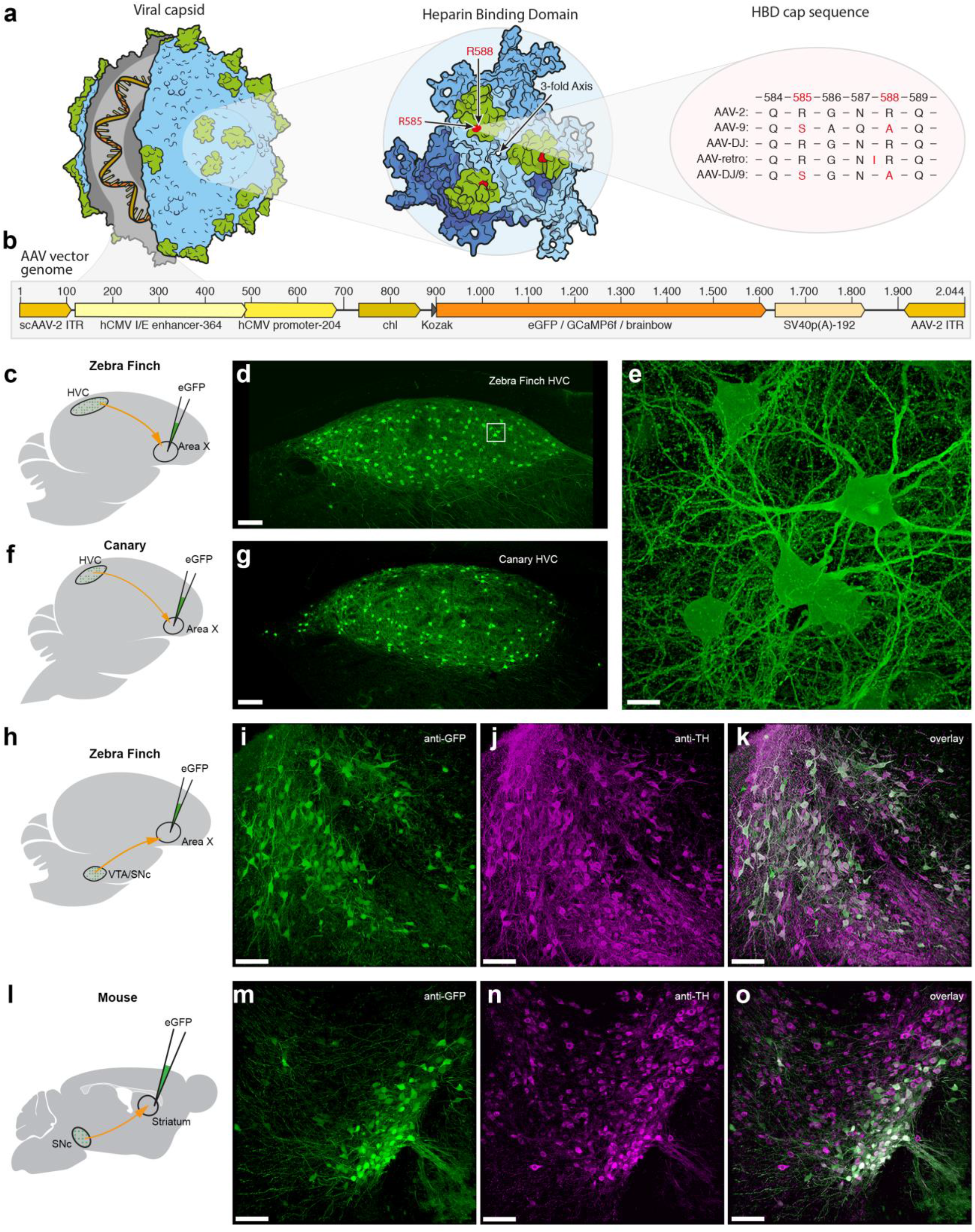
AAV capsid and genome structure for efficient retrograde access to songbird and mouse projection and dopaminergic neurons. **a**, Illustration of the AAV capsid, showing exposed surface proteins responsible for binding to the heparin sulfate proteoglycans (HSPG) receptor, thought to be important for cellular entry of the viral vector. Mutations of two crucial arginines in the heparin binding domain (HBD, highlighted in green) cap sequence at positions 585 and 588 (highlighted in red) reduce heparin binding affinity and potentially improve retrograde transport. Changes are shown for four serotypes that have been used for or possess potential for retrograde transduction. **b**, AAV vector genomes encoding diverse transgenes (orange region) including fluorescent proteins and the calcium indicator GCaMP6f. Numbers on top indicate the base pair length and relative position of encoded elements. Note that the actual size of the region encoding the transgene (orange) varies depending on the respective protein. Brainbow stands for 3 separate AAV vector genomes encoding either eGFP (enhanced green fluorescent protein), eCFP (enhanced cyan fluorescent protein), or mRuby3 (red fluorescent protein). **c**, **f**, **h**, **l**, Schematics of the virus injection sites (green) and the neuronal projections analyzed (red). **d**, **g**, Native fluorescence signal in retrogradely labeled HVC_X_ neurons 7 days post virus delivery in zebra finch and in canary, respectively. **e**, Higher magnification z-stack of the boxed region in d. Dopaminergic projection neurons in zebra finch (**i-k**) and mouse (**m-o**) retrogradely labeled with our eGFP-construct, after immunostaining for anti-GFP (**i**, **m**) and anti-Tyrosine hydroxylase (**j**, **n**). Scale bars 100 μm except e, 10 μm.

## Results

### Fast and efficient retrograde transduction of songbird projection neurons

First, we examined the transgene expression kinetics and stability of our viral construct. We assessed expression performance in the corticostriatal connection from nucleus HVC (proper name) to Area X. HVC, the songbird vocal premotor cortex analog, is a major hub during vocal learning, integrating information from auditory^18^, dopaminergic^15^, and premotor^19^ afferents. HVC includes three major neuron populations: local interneurons, HVC_RA_ neurons that project to the robust nucleus of the arcopallium (RA), which innervates vocal and respiratory motor neurons, and HVC_X_ neurons that project to Area X, the basal ganglia part of a cortico-basal ganglia-thalamic loop involved in song learning^20^. HVC_X_ neurons are analogous to mammalian corticostriatal projection neurons, which are involved in motor learning in mammals^21^.

We found robust native fluorescence of eGFP in HVC_X_ neurons as early as 3 days post-injection of scAAV-2-DJ/9-hCMV-I/E-chI-eGFP-p(A) into Area X (Fig. 2a,b). Abundant eGFP-filled neurons could be seen within HVC throughout four different time points ranging from 3 to 28 days after injection (Fig. 2a-h). Three days after injection, we detected eGFP labeling among 59% (15.56 x 10^3^ somata/mm^3^, n=6 hemispheres) of HVC_X_ somata as compared to retrograde labeling with Cholera Toxin B (CTB, 26.48 x 10^3^ somata/mm^3^, n=8 hemispheres; Fig 2i,j), a commonly used and highly efficient retrograde tracer in zebra finches^22^. Already 7 days post-injection, eGFP labeling density (23.52 x 10^3^ somata/mm^3^, n=8 hemispheres; Fig. 2c,j) was comparable with CTB labeling density, hinting at a strikingly high transduction efficiency. Longer eGFP expression times of 14 and 28 days did not further increase eGFP labeling densities (25.28 x 10^3^ and 23.66 x 10^3^ somata/mm^3^ respectively, n=8 hemispheres per time point; Fig. 2e,g,j). These results suggest that the peak of native fluorescence of eGFP expression is reached already 7 days post-delivery and that there is no obvious decay of fluorescence or neurotoxicity for at least 4 weeks after injection.

**Fig. 2.**
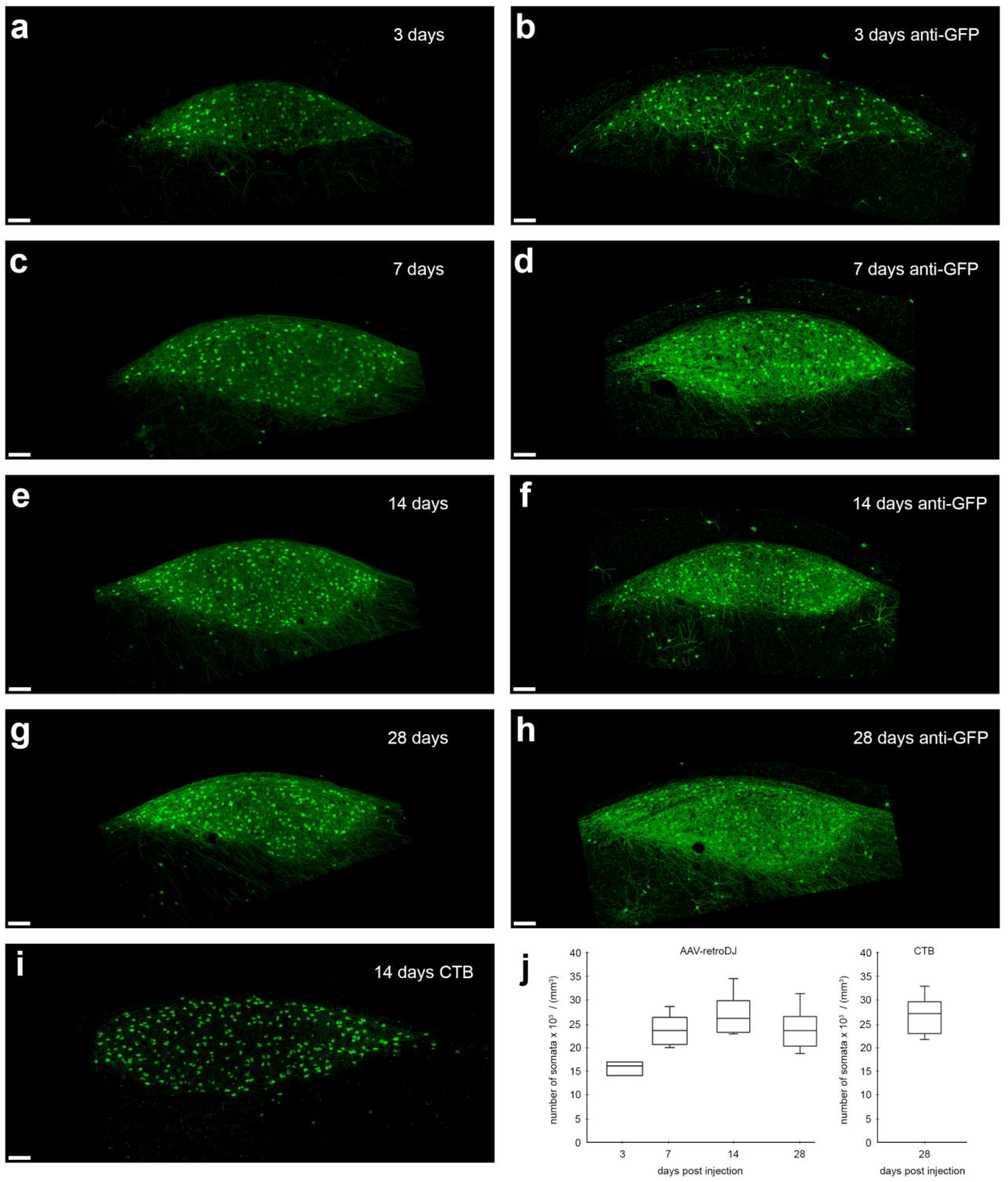
Retrograde transduction kinetics in zebra finch HVC_X_ neurons. We find strong retrograde expression of eGFP in Area-X projecting (HVC_X_) neurons as early as 3 days after injection of scAAV-2-DJ/9-hCMV-I/E-chI-eGFP-p(A) into Area X. Confocal images show native eGFP signal postdelivery after **a**, 3 days; **c**, 7 days; **e**, 14 days; and **g**, 28 days; and **b**, **d**, **f**, **h**, their respective anti-GFP immunostainings. **i**, Comparative image of retrogradely labeled HVC_X_ neurons 14 days after injections of cholera toxin B (CTB, a retrograde neural tracer) into Area X. Note the strong eGFP signal in neurites of virally labeled cells as early as 3 days as compared to the neural tracer. **j**, Number of retrogradely labeled somata of HVC_X_ neurons after 3, 7, 14, and 28 days post-delivery into Area X (based on native eGFP signal) compared to 14 days labeling with the retrograde tracer. Scale bars 100 μm.

### Transduction success rate and specificity of retrograde projection-neuron labeling in songbirds

We assessed the transduction success rate of our eGFP-construct, and found consistent transduction of HVC_X_ neurons in 24 out of 24 single-injected hemispheres (n=12 birds) after a minimum of 7 days after virus injection into Area X. For comparison, previous studies applying AAV vectors for retrograde transduction in songbirds have reported overall successful transduction in as little as 18% of virus injections (one brain pathway showing retrograde eGFP expression in 2 out of 11 injected hemispheres)^9^. To date, we have not encountered a single injection of our eGFP-construct without successful retrograde transduction of projection neurons after 7 days.

We further assessed the specificity of retrograde labeling. HVC and Area X are anatomically well separated, which virtually excludes the possibility that viral particles passively diffuse from Area X to HVC. In addition, the fact that this connection is unidirectional makes it safe to assume that all labeled neurites in HVC belong to HVC_X_ neurons. Nonetheless, some studies report the possibility of trans-synaptic spread of AAV vectors^23,24^. To assess whether all labeled neurons are indeed HVC_X_ neurons, and to exclude any potential mislabeling, we examined the morphological characteristics of eGFP-labeled HVC neurons.

The three main HVC neuron classes are morphologically well described and can be separated based on dendritic spine densities^25^. HVC interneurons are mainly aspiny, HVC_RA_ projection neurons are moderately spiny (0.21 ± 0.07 spines/μm), and HVC_X_ neurons have the highest dendritic spine densities (0.70 ± 0.13 spines/μm). We analyzed 17 randomly chosen dendritic branches of various lengths (total branch length of 396 μm) across n=4 adult male zebra finches (>7 days after injection). A subset of histological brain sections was further subjected to anti-GFP immunolabeling to capture any potentially low-transgene expressing cells. Analyzed dendritic branch fragments had dendritic spine densities ranging from 0.47 to 1.75 spines/μm (average 1.03 ± 0.35 spines/μm, which compares favorably with the previously reported 0.70 ± 0.13 spines/μm^25^), indicating that eGFP-expressing cells are most likely all HVC_X_ neurons. Albeit some overlap in spine density between X- and RA-projecting cells, we did not encounter a single cell that was obviously either aspiny (interneuron) or had a very low spine density (HVC_RA_ neurons).

To further exclude the possibility of unspecific transduction of non-neuronal cell types such as astrocytes, we performed anti-neuronal protein HuC/HuD counterstains, a common neuronal marker. We found that all examined eGFP-labeled cells were also labeled with HuC/HuD antibody (n=4 sections), confirming their neuronal identity (Fig. 3).

**Fig. 3.**
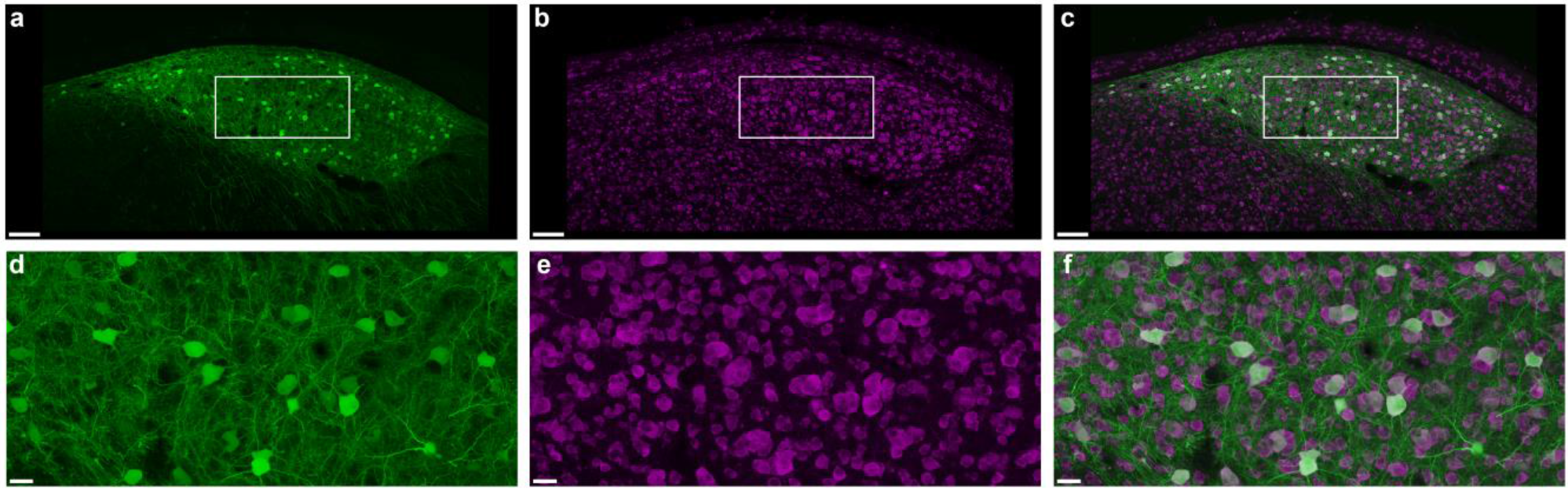
Retrograde expression is restricted to neuronal cell types. **a**, Confocal microscopy image of zebra finch HVC with retrogradely labeled HVC_X_ neurons. **b**, Immunostaining with anti-HuC/HuD, a commonly used antibody against neurons. **c**, Overlay of eGFP signal (green) and HuC/HuD signal (magenta), showing no HVC_X_ neurons unlabeled for HuC/HuD. **d**, **e**, **f**, Higher magnification of the boxed regions in a, b, c. Scale bars a, b, c, 100 μm; d, e, f, 20 μm.

### Visualization of morphological details in retrogradely labeled projection neurons

The weak transgene expression produced by previous retrograde vectors for songbirds typically results in fluorescent protein-labeling that is often limited to somata and small extensions of proximal neurites^9,13^. Using our new eGFP-construct thus opens new possibilities for highly detailed morphological analysis of songbird projection neurons. In fact, the high expression levels of eGFP in HVC_X_ neurons 7 days post-delivery allowed a remarkable visualization of dendritic morphological details. Adjacent to imaged dendrites, we found clearly distinguishable dendritic spines throughout HVC (Fig. 1e), at sufficient detail to distinguish and trace different spine types (Fig. 4).

**Fig. 4.**
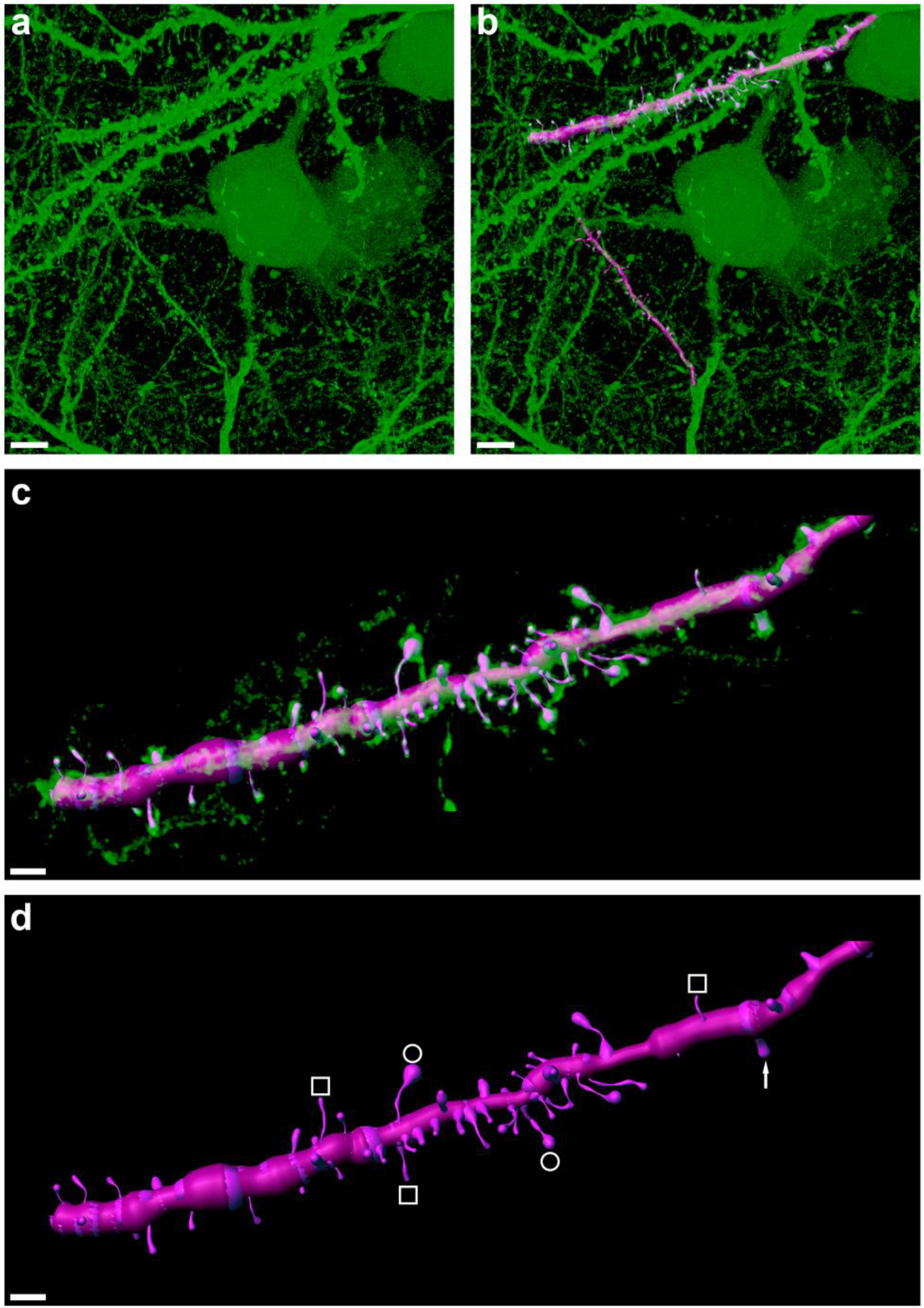
Strong eGFP expression permissive for reconstructing dendritic spine morphologies. **a**, High magnification confocal image stack of retrogradely labeled HVC_X_ neurons showing strong expression of eGFP in dendritic fragments. **b**, Digital 3D-reconstruction (magenta overlay) of two exemplary dendritic fragments. **c**, Isolated and zoomed view of the upper dendritic fragment shown in b, revealing high morphological detail. **d**, In the semi-automatically reconstructed dendritic fragment, various spine morphologies can be recognized including philopodia (white squares), mushroom heads (white circles), and stubby spines (white arrow). Scale bars a, b, 5 μm; c, d, 2 μm.

The extensive eGFP labeling makes our construct a promising candidate for *in vivo* imaging of morphological plasticity. To date, *in vivo* imaging of dendritic spine plasticity in songbirds has been achieved exclusively via lentivirus-mediated fluorescence labeling^12,26^, a tool that lacks retrograde transduction capabilities. Furthermore, the strong eGFP expression we demonstrate is highly beneficial for tissue processing techniques that require a strong initial fluorophore expression, including large-volume expansion (lattice) light-sheet microscopy (ExLSM^27^, ExLLSM^28^) and whole-brain tissue clearing^29^.

### Retrograde expression in dopaminergic projection neurons

Similarly to the mammalian basal ganglia, Area X is densely innervated by dopaminergic neurons of the ventral tegmental area-substantia nigra pars compacta complex (VTA/SNc), forming a continuous group of cells with diverse projection targets and innervation^30^. In humans, the comparable dopaminergic system is crucially affected by neurodegenerative disorders such as Parkinson’s disease^31^. Songbird VTA/SNc_X_ neurons (traditionally referred to as VTA_X_ neurons) play crucial roles in vocal learning, as they are responsible for encoding reward prediction errors associated with singing^32^.

In birds injected with our construct into Area X, we observed good retrograde eGFP labeling of VTA/SNc_X_ neurons. To identify the types of the labeled neurons, we immunolabeled histological sections for tyrosine hydroxylase (TH), which is a good marker for dopamine and has been shown to label the majority of VTA/SNc_X_ neurons (estimates ranging from 88 to 95% of VTA/SNc_X_ neurons)^11,14,33^. We found that the majority of examined VTA/SNc_X_ neurons were indeed positively labeled for TH (n=4 birds, Fig. 1h-k), suggesting that these neurons are dopaminergic.

The strength of retrograde labeling of dopaminergic VTA/SNc projection neurons in zebra finches led us to investigate whether high labeling efficiency can also be observed in rodents. One of the most effective retrograde vectors in rodents, AAV-retro, only poorly transduces mouse dopaminergic projection neurons in the SNc^34^. Coincidently, we tested AAV-retro and found this construct to also be non-functional in zebra finches (n=4 adult male zebra finches, single injections into Area X in both hemispheres of the construct scAAV-2-retro-hCMV-I/E-chI-eGFP-p(A), with post-delivery incubation times of 6 weeks). When we injected our eGFP-construct into the striatum of mice, a major projection target of the SNc, we found successful labeling of dopaminergic SNc projection neurons (confirmed with TH-immunolabeling, n=5 mice, Fig. 1l-o).

### Retrograde expression in canary projection neurons

The strong expression and retrograde transport of our new viral construct in both zebra finches and mice hints towards a broad species applicability of this vector. We therefore sought to test our construct on canaries. Canaries (*Serinus canaria*) are seasonal breading songbirds that produce highly complex and flexible songs with extremely fast syllable repetitions, making this species a valuable model system for complex motor skill learning^35^. Additionally, canaries are a popular model for sex hormone-induced neuroplasticity^36^. Seasonal changes in singing behavior in this species are correlated with sex hormone fluctuations and striking neural changes. To date, the application of viral vectors in the canary brain has been limited to the use of lentiviruses^37^, which do not retrogradely transduce projection neurons. Injections of our eGFP-construct into canary Area X resulted in strong eGFP expression in HVC_X_ neurons 7 days after injection, confirming the rapid expression times observed in zebra finches (Fig. 1f,g).

### Brainbow labeling of projection neuron target circuits

One common problem associated with studying the structure of local neural circuits is excessive labeling density, which can hinder the visual separation of adjacent neurons and their respective dendrites. One tool that can potentially overcome this problem is brainbow labeling^38^. Brainbow labeling color-codes neurons via the relative expression ratios of diverse fluorescent proteins. One brainbow technique achieves diverse labeling colors via the selective uptake ratios of diverse viral particle types. This technique involves simply mixing and co-injecting three individual vector types encoding different chemical tags or fluorophores^39^ (Fig. 5a). Brainbow labeling in any of its variants has not yet been applied in songbirds.

**Fig. 5.**
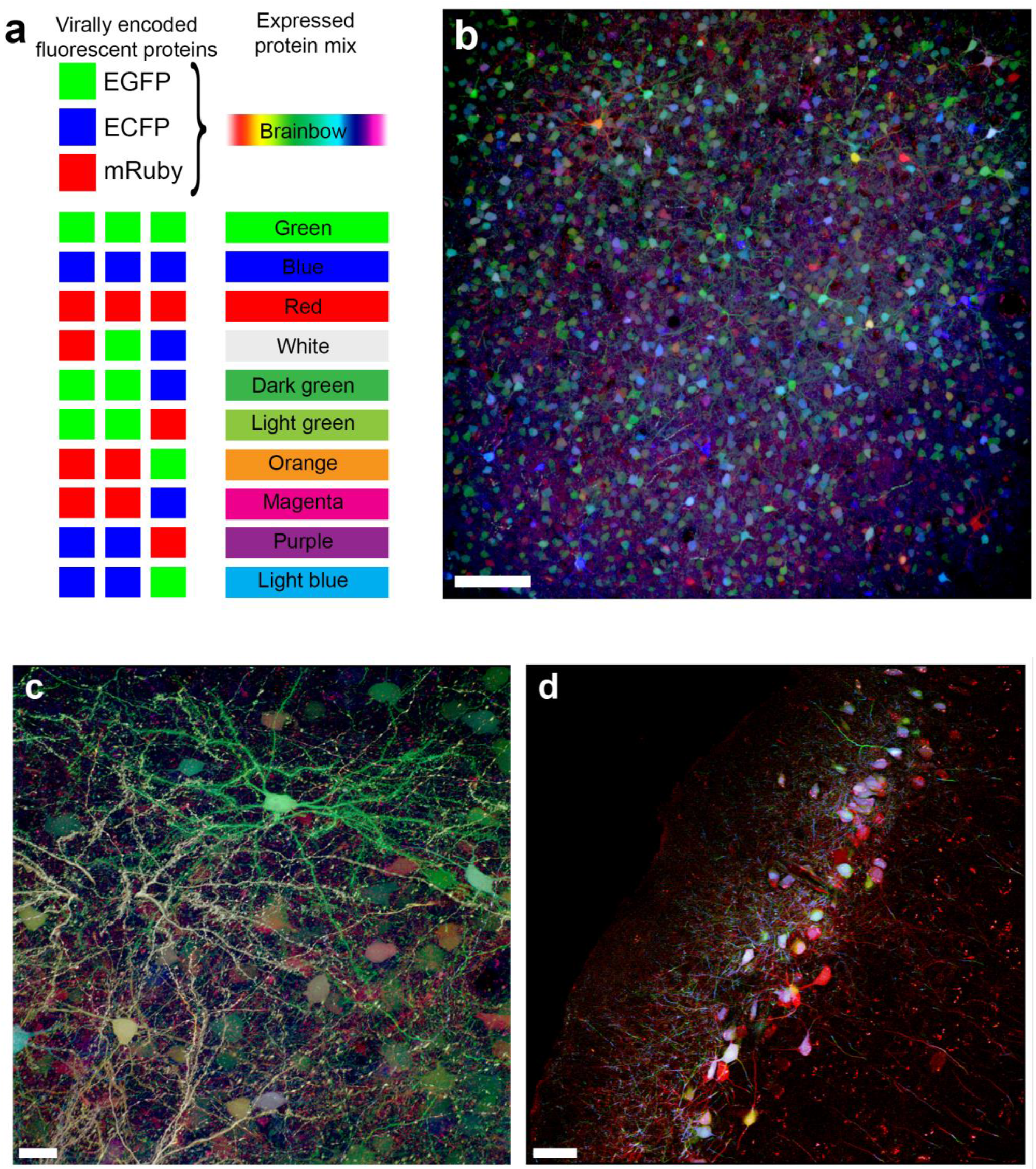
Brainbow labeling in zebra finch and mouse. **a**, AAV-mediated brainbow labeling is achieved by injecting a mixture of three individual vectors encoding either eGFP, eCFP or mRuby3. Each neuron ends up displaying a distinct color because of variable transduction of the three fluorescent proteins, largely depending on the number of vectors per cell per fluorophore. **b**, Brainbow labeling of locally transduced Area X neurons allows for recognition of individual neurons in a densely labeled z-stack. **c**, High magnification confocal imaging reveals consistent color labeling in dendritic fragments of Area X neurons, including spines. **d**, Retrograde brainbow labeling of hippocampal projection neurons in mouse. Scale bars b, 100 μm; c, 20 μm; d, 50 μm.

In addition to the eGFP-expressing vector, we produced two additional vectors expressing spectrally distinct fluorescent proteins, enhanced cyan fluorescent protein (eCFP) and red fluorescent protein (mRuby3). Next, we injected the three aforementioned constructs in a ratio of 1:1:1 into Area X of adult male zebra finches. Local expression within Area X produced spectrally diverse neuron labeling (Fig. 5b) with consistent and dense fluorophore expression patterns throughout neurons including morphological fine structures such as spines (Fig. 5c). Although abundant brainbow labeling strategies exist for mice, we are not aware of any AAV-based retrograde labeling strategy which does not involve Cre-Lox recombination. We injected our eGFP-, eCFP-, and mRuby3-constructs in a ratio of 1:1:1 into the murine area CA1 of the dorsal hippocampus (n=4). The injection of these three constructs produced spectrally diverse labeling of projection neurons in the mouse entorhinal cortex (Fig. 5d).

### Retrograde expression of calcium sensors

In light of the obvious benefits of our construct for visualization of projection neuron circuits, we sought to investigate whether the expression of other genetic cargo could be mediated equally well. The packaging capacity of natural AAV serotypes is limited to a genome size of about 4.7 kb. Nevertheless, the absolute packaging limit of AAV vectors has been challenged and one recent study found a brick-wall limit of 5.2 kb for AAV-8^40^. Because the maximum packaging size of an AAV likely also depends on the exact protein composition of the nucleocapsid, we decided to produce the same viral construct for GCaMP6f, with a total genome size of 5.286 kb, which is just over the reported maximum. Production was successful and yielded a physical viral titer of 5.9 x 10^12^ vg/ml, which seemed promising to test for *in vivo* calcium imaging.

GCaMP is a genetically encoded calcium (Ca^2+^) sensor^41^ consisting of a circular GFP, calmodulin (CaM) and a peptide chain (M13), which in its natural conformation shows only poor fluorescence. In the presence of Ca^2+^, CaM undergoes a structural change that entails a rapid increase in fluorescence. GCaMP6f was engineered for fast fluorescence dynamics and high Ca^2+^ sensitivity, resulting in reliable single-spike detection at 50-75 ms inter-spike intervals^42^.

We injected our GCaMP6f-construct into the RA of Bengalese finches (*Lonchura striata var. domestica*), a further songbird species commonly used in birdsong research. We used a custom-built miniaturized fluorescence microscope (see Methods) to image neuronal activity *in vivo* under isoflurane anesthesia. We imaged spontaneous activity in HVC_RA_ neurons (Fig. 6, Supplementary Video 1), an HVC projection neuron population that generates precise temporal sequences^43^ during song production. In a field of view of (262.5 μm)^2^, we were able to detect 48 spiking events in 8 distinct HVC_RA_ projection neurons over a time course of 100 s.

**Fig. 6.**
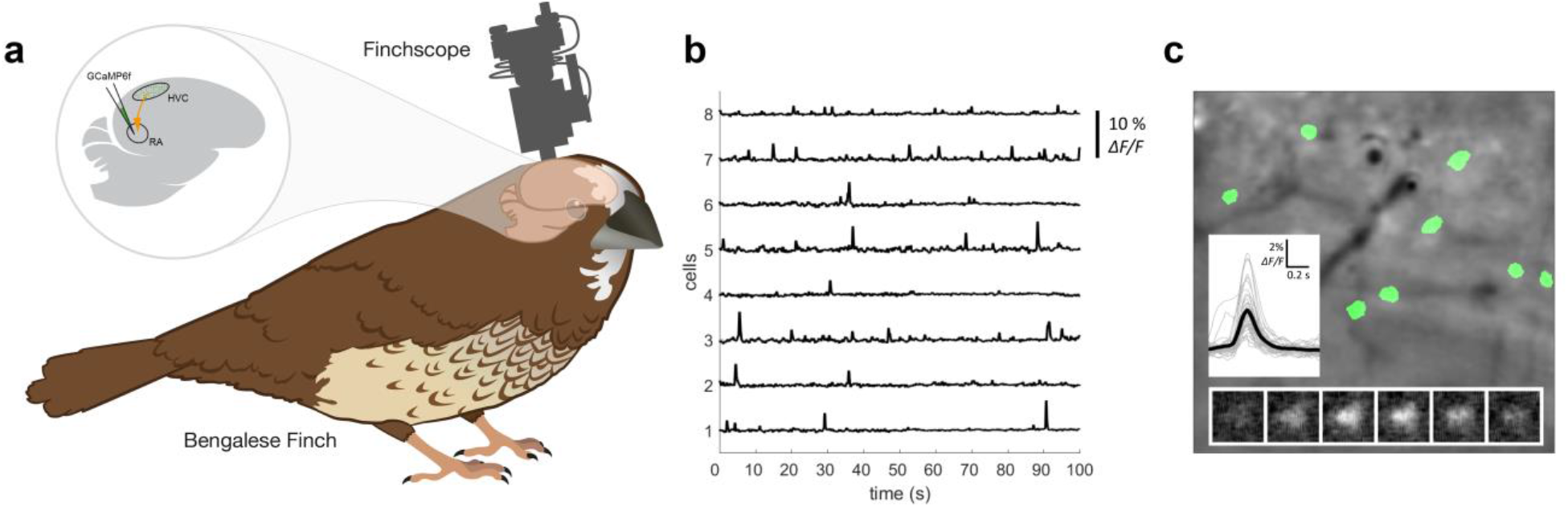
*In vivo* calcium imaging in retrogradely labeled Bengalese finch RA-projecting (HVC_RA_) neurons. **a**, Schematic of the scAAV-2-DJ/9-hCMV-I/E-chI-GCaMP6f -p(A) injection site and miniature microscope (finchscope) attachment in Bengalese finches. **b**, Fluorescence intensity traces (ΔF/F) for eight individual cells expressing GCaMP6f. All cells show clear spontaneous spike-like activity as indicated by a sudden increase in signal intensity. **c**, Single frame of *in vivo* calcium fluorescence movie under isoflurane anesthesia. Regions tracked over time are indicated in green. Bottom inset shows fluorescence signal of one cell over a period of 0.5 s (100 ms between snapshots). Left inset shows mean calcium transient (n=43) in black and individual transients in grey (windows containing multiple events were excluded for visual clarity).

## Discussion

We present a new viral construct for retrograde delivery of genetic cargo to projection neuron circuits in songbirds, with fast and robust transgene expression and high transduction efficiency. Our construct, scAAV-2-DJ/9-hCMV-I/E-chI-transgene-p(A), is suitable for studying the detailed connectivity and function of songbird corticostriatal and vocal motor pathways essential for vocal learning. Our construct also provides reliable access to probe the structure of cortical and dopaminergic projection neuron circuits in both songbirds and mice. These findings indicate the applicability of our new vector in both species and circuits that appear to be resistant to retrograde targeting with AAV vectors.

The current lack of reliable tools to target specific neuron populations in songbirds has been a possible contributor to the underrepresentation of songbirds as model species in medical and applied research. For targeted manipulations of projection neuron populations in the zebra finch, one serotype, AAV-9, has previously been used^9–11,13^. However, these vectors entail significant drawbacks, such as low transduction success rates, prolonged incubations times, and sparse and weak transgene expression, all of which limit their application. These limitations were not remedied by the use of the presumably stronger-expressing self-complementary variant of AAV-9^9,13^, nor by exploiting the cre-flex system with the cre-recombinase vector injected at the projection target^10,11,13^. Unlike in songbirds, in mice, the cre-flex system is very efficient and extremely low levels of cre are able to drive expression of otherwise silenced transcripts^44^.

Despite previous limitations as retrograde tools for the songbird brain, AAV vectors still seemed worth exploration thanks to their extensive capsid variety, with more than 100 existing capsid variants. The inability of certain viral vectors to infect songbird tissue is likely owed to diverse factors. Such factors possibly include incompatibility of co-receptors for cellular access of the viral vector, as well as differences in promoter sequences and intracellular physiology, both of which can impair transduction. A further potential limiting factor is a substantially different immune response in birds compared to mammals. Commonly, immune defense systems pose the first barrier when exploring viral tools. Accordingly, the low transduction success rate of AAV-9 vectors hints towards a possible strong immune response in zebra finches against this serotype. This hypothesis led us to explore serotypes focusing on low immune response, which is a trait of vectors engineered for human gene therapy. Human gene therapy vectors are required to have high transduction efficiency and low immunogenicity, both of which were highly desirable characteristics for the new songbird viral tool we were pursuing.

AAV serotypes have previously been selected in order to yield high efficiency and low immunogenicity^17^. The single prevailing capsid, the AAV2, 8 & 9 chimera termed AAV-DJ, showed superior transduction characteristics, which motivated us to test this capsid. We found that the self-complementary variant of AAV-DJ shows great local expression in songbird brain tissue, with great transduction characteristics, but unfortunately shows no retrograde transduction. Although showing some potential for retrograde access to projection neurons^34,45,46^, AAV vectors are traditionally not used for retrograde studies. One notable exception, the AAV-retro construct, has demonstrated a 10-fold higher retrograde transduction efficiency than AAV-DJ^34^. This motivated us to test AAV-retro, but we found this construct to be non-functional in zebra finches. We thus carefully examined the changes introduced in AAV-retro, which presumably led to increased efficacy of retrograde access. The critical changes appear to interact with the heparin sulfate proteoglycans (HSPG) binding domain (HBD)^47^. The 10-mer insert between positions 587 and 588 (Fig. 1a) probably both disrupts HBD’s functionality, as demonstrated by the reduced heparin binding affinity of AAV-retro, and creates a new binding surface, which might improve vesicular trafficking or nuclear entry of viral particles^34^. Interestingly, the characteristics of the HBD were also carefully examined by the creators of the AAV-DJ construct^17^. One of their negative controls produced for comparison to AAV-DJ, AAV-DJ/9, included two point mutations in the HBD that disrupt heparin binding. These two point mutations seemed to reduce transduction efficiency of AAV-DJ/9 in comparison to AAV-DJ, but surprisingly also induced faster transduction kinetics. Faster transduction kinetics was a highly desirable characteristic for the new songbird viral tool we were pursuing, which peaked our interest in the AAV-DJ/9 capsid variant. Coincidentally, the point mutations of the HBD in AAV-DJ/9 also rendered this serotype closer to its parental serotype AAV-9^17^ (Fig. 1a), which has been shown to have some potential for retrograde access in songbirds^10–12,14^.

The exact composition of our construct combines many features that most likely all contribute constitutively to the highly efficient transgene expression and retrograde transport (Fig. 1a,b). Clearly, the AAV-DJ/9 capsid structure of the HBD plays a major role for the viral access through axon terminals and the fast transduction kinetics. Nonetheless, this feature alone does not guarantee strong retrograde expression. The viral vector also needs to be transported by the transduced cells lysosomal system into the nucleus, where the genetic cargo has to be translated into messenger RNA (mRNA). The process of conversion into mRNA in the nucleus can be drastically accelerated by packaging two complementary copies of single stranded DNA genomes in a self-complementary AAV vector instead of single stranded DNA genomes. However, self-complementary AAV vectors have also been shown to induce a stronger immune response^48^. When applying a self-complementary AAV9-based vector (scAAV9) for retrograde access in songbirds, the self-complementary genome seems to neither have improved transduction success rates, transduction efficiency, nor expression kinetics^9^. The disappointing results obtained with scAAV9 might, however, be partially explained by the chosen promoter. Although a wide range of promoters have been applied to songbird brain tissue, we are not aware of a promoter that stands out in projection neurons. The scAAV9 vector has been employed in songbirds using a CBh promoter^9^, a hybrid variant of the chicken beta actin (CBa) promoter. Nonetheless, CBh supposedly ensures strong expression through self-complementary vector mediated transduction in neuron types that are also affected by CMV promoters^49^. Given the poor transduction efficiency of scAAV-9-CBh, it could be that the hCMV immediate-early enhancer (hCMV-I/E) and/or the chimeric intron (chI) of our self-complementary construct play crucial roles for retrograde transduction in projection neurons. The introduced intron might contribute to the improved transduction efficiency by regulating splicing of mRNA within the nucleus, which has been shown to improve nuclear export^50^. Further investigations would be necessary to fully elucidate the exact contributions of individual components to retrograde transport and transduction efficiency and kinetics.

In this work we present a new viral construct for excellent access to specific projection neuron populations in songbirds. We demonstrated the suitability of our new construct for *in vivo* imaging of calcium activity (Fig. 6), and for detailed morphological analysis based on extensive axonal and dendritic labeling, using both single (Fig. 4) and multi-color approaches (Fig. 5). The high transduction success rates and great transduction efficiency hint at the potential for this tool to make a significant contribution to animal welfare by reducing the number of experimental animals required in future studies, as stated by the 3R principles. Moreover, our new tool will likely open new avenues of investigation into the structure and function of projection neuron populations and allow the study of the brain circuits underlying vocal learning in unprecedented detail, including relevant dopaminergic inputs during song development.

## Acknowledgments

This work was supported by European Union’s Horizon 2020 Marie Skłodowska-Curie grant Nr. 750055 (D.D.), ETH grants ETH-42 15-1 and ETH-20 19-01 (R.H. and B.G.), SNSF Sinergia grant CRSII5-173721 (B.G.), and Swiss Data Science Center Grant C17-18 (B.G.). Part of the imaging was performed at the Center for Microscopy and Image Analysis, University of Zurich. Some figures were generated with the help of the Scientific Illustration and Visual Communication (SIVIC) center of the University of Zurich. The AAV-DJ/9 helper plasmid was a kind gift from Mark A. Kay (Department of Pediatrics & Genetics, Stanford University). The plasmid pBV1 was a kind gift from Bernd Vogt (Institute of Virology, University of Zurich). We thank Veronika Bednarova and Benedikt Grothe (Max-Planck Institute of Neurobiology) for preliminary experiments in mice.

## Author contributions

DD, RH and FD conceived of the study. JC, MR, and DD designed viral constructs. JC and MR produced all vectors. DD, FD, MDR, RT, CM, and RB performed experiments. DD, FD, and RK contributed to confocal imaging. All authors contributed to image interpretation and data analysis. MG, KO, RH, BG, RK, and JC provided equipment, reagents and materials. DD prepared all figures. DD and MDR wrote the manuscript. All authors contributed to manuscript writing or revision and approved of the final version.

## Supplementary Files

**Supplementary Video 1 | *In vivo* calcium imaging in retrogradely labeled HVC_RA_ neurons in the Bengalese finch.** Video showing retrogradely transduced HVC_RA_ neurons expressing GCaMP6f (bright spots). Eight cells show clear activity as indicated by the change in local signal intensity (compare also to Fig. 6).

## Methods

### Design of self-complementary adeno-associated virus vector plasmids

Self-complementary adeno-associated virus (AAV) vector plasmids (pscAAV) were constructed as previously described^1,2^. Briefly, the terminal resolution site (trs) and the packaging signal (D-sequence) from psub-2-CBA-WPRE^3,4^ were deleted by *Bal* I restriction digestion within the AAV serotype 2 (AAV-2) 5’ inverted terminal repeat (5’-ITR) resulting in the plasmid pscAAV-2-Δ3’-ITR. Subsequently, the multiple cloning site (MCS) of pBluescript II SK (+) (Stratagene) together with the AAV-2 3’-ITR and simian virus 40 late polyadenylation signal (SV40p(A)) containing fragment of psub-2-CMV-WPRE^3^ were inserted into pscAAV-2-Δ3’-ITR, resulting in the plasmid pscAAV-2-MCS-SV40p(A). In the AAV vector plasmids used here (Fig. 1b), the human cytomegalovirus (hCMV) promoter/immediate-early enhancer (IE) of peGFP-N1 (Clontech) and the chimeric intron (chI) of pSI (Promega) were inserted into pscAAV-2-MCS-SV40p(A) resulting in pscAAV-2-hCMV-chI-SV40p(A).

The eGFP open reading frame (ORF) was amplified by PCR using pscAAV-2-hCMV-chI-floxedeGFP as the template DNA and primers 5’-ATACTAGTGCCACCATGGTGAGCAAGGGCG-3’ (forward) and 5’-TTGCGCGGCCGCTTACTTGTACAGCTCGTCCATG3’ (reverse). Amplicons were Spe I/Not I restriction digested and inserted into the Spe I/Not I restriction digested pscAAV-2-hCMV-chI-floxedeGFP to generate pscAAV-2-hCMV-chI-eGFP (eGFP vector plasmid). For construction of the mRuby3 vector plasmid (pscAAV-2-hCMV-chI-mRuby3-SV40p(A)), the mRuby3 ORF was amplified by PCR using Addgene #85146 as the template DNA and primers 5’-CATTACTAGTGTTTAAACACTCGAGGCTAGCGCCACCATGGTGTCTAAGG-3’ (forward) and 5’-TAGGCGCGCCTACGTACAATTGGGTACCTTACTTGTACAGCTCGTCCATG-3’ (reverse). The resulting PCR product was cut with SpeI and BsrGI and inserted into the *Spe* I and *BsrG* I sites of the eGFP vector plasmid.

For construction of the eCFP vector plasmid (pscAAV-2-hCMV-chI-eCFP-SV40p(A)), the eCFP ORF was isolated from plasmid pBV1 as *Nhe* I/*Sac* II fragment and inserted into the *Sac* II/*Spe* I opened eGFP vector plasmid.

For construction of the GCaMP6f vector plasmid (pscAAV-2-hCMV-chI-GCaMP6f-SV40p(A)), the GCaMP6f ORF was isolated by Bgl II/BstB I restriction digest from pssAAV-2-hSyn1-chI-GCaMP6f-WPRE-SV40p(A) (N-terminal part of GCaMP6f) and by BstBI/BssH II(blunt) restriction digest from pssAAV-2-hEF1a-dlox-GCaMP6f(rev)-dlox-WPRE-bGHp(A) (C-terminal part of GCaMP6f) and inserted into the BsrG I(blunt)/Bgl II restriction digested pscAAV-2-hCMV-chI-Lck_eGFP-SV40p(A) to generate pscAAV-2-hCMV-chI-GCaMP6f-SV40p(A) (GCaMP6f vector plasmid).

The identity of all constructs was confirmed by Sanger DNA sequencing and restriction endonuclease analyses.

Sequences of all viral vectors and their corresponding plasmids can be found in the repository of the Viral Vector Facility of the University of Zurich and ETH Zurich (https://www.vvf.uzh.ch/en.html).

### Production, purification, and quantification of self-complementary (sc) AAV vectors

Self-complementary (sc) AAV vectors were produced and purified as previously described^4,5^. Briefly, human embryonic kidney (HEK) 293 cells^6^ expressing the simian virus (SV) large T-antigen^7^ (293T) were transfected by polyethylenimine (PEI)-mediated cotransfection of AAV vector plasmids (providing the to-be packaged AAV vector genome, see above), the AAV helper plasmid pAAV-DJ/9; pAAV-DJ/9 providing the AAV serotype 2 rep proteins and the cap proteins of AAV-DJ/9) and adenovirus (AV) helper plasmids pBS-E2A-VA-E4^3^ (providing the AV helper functions) in a 1:1:1 molar ratio.

At 120 to 168 hours post-transfection, HEK 293T cells were collected and separated from their supernatant by low-speed centrifugation. AAV vectors released into the supernatant were PEG-precipitated over night at 4 °C by adding a solution of polyethylenglycol 8000 (8% v/v in 0.5 M NaCl), and completed by low-speed centrifugation. Cleared supernatant was discarded and the pelleted AAV vectors resuspended in AAV resuspension buffer (150 mM NaCl, 50 mM Tris-HCl, pH 8.5). HEK 293T cells were resuspended in AAV resuspension buffer and lysed by Bertin’s Precellys Evolution homogenizer in combination with 7 ml soft tissue homogenizing CK14 tubes (Bertin). The crude cell lysate was DENARASE (c-LEcta GmbH) treated (150 U/ml, 90 to 120 minutes at 37 °C) and cleared by centrifugation (10 minutes at 17.000 g/4 °C). The PEG-precipitated (1 hour at 3500 g/4 °C) AAV vectors were combined with the cleared cell lysate and subjected to discontinuous density iodixanol (OptiPrep™, Axis-Shield) gradient (isopycnic) ultracentrifugation (2 hours 15 minutes at 365’929 g/15 °C). Subsequently, the iodixanol was removed from the AAV vector containing fraction by 3 rounds of diafiltration using Vivaspin 20 ultrafiltration devices (100’000 MWCO, PES membrane, Sartorius) and 1x phosphate buffered saline (PBS) supplemented with 1 mM MgCl_2_ and 2.5 mM KCl according to the manufacturer’s instructions. The AAV vectors were stored aliquoted at −80 °C.

Encapsidated viral vector genomes (vg) were quantified using the Qubit™ 3.0 fluorometer in combination with the Qubit™ dsDNA HS Assay Kit (both Life Technologies). Briefly, 5 μl of undiluted (or 1:10 diluted) AAV vectors were prepared in duplicate. Untreated and heat-denaturated (5 minutes at 95 °C) samples were quantified according to the manufacturer’s instructions. Intraviral (encapsidated) vector genome concentrations (vg/ml) were calculated by subtracting the extraviral (non-encapsidated; untreated sample) from the total intra- and extraviral (encapsidated and non-encapsidated; heat-denatured sample). All AAV vectors used in this study had vector genome concentrations between 3.8 x 10^12^ vg/ml and 7.4 x 10^12^ vg/ml.

Identity of encapsidated genomes were verified and confirmed by Sanger DNA-sequencing of amplicons produced from genomic AAV vector DNA templates (identity check).

### Animals

Zebra finches were obtained from breeding colonies in Zurich, Switzerland, or Seewiesen, Germany; canaries from breeding colonies in Seewiesen, Germany; and Bengalese finches from breeding colonies in the University of Tokyo, Japan. C57BL/6 mice were obtained from Charles River Germany and housed in the LASC animal husbandry at the University of Zurich.

Animal handling and all experimental procedures were conducted following the ethical principles and guidelines for animal experiments of Switzerland/Germany/Japan.

### Surgical procedure for viral vector delivery

Birds were deeply anesthetized using an orally administered mixture of oxygen with 1-3% isoflurane gas before being placed in a custom stereotaxic apparatus. After applying a topical anesthetic, a vertical incision was made in the skin over the skull, and a small craniotomy was performed at predetermined distances from the anatomical landmark lambda. For bi-lateral injections into Area X the following stereotaxic coordinates where used relative to lambda: anterior-posterior-axis (AP) +5.4 mm, medial-lateral-axis (ML) 1.6 mm, and dorsal-ventral-axis (DV) −3.2 mm, with an approximately 85 degree earbar-beak angle. For bi-lateral injections into RA of Bengalese finches we used the following coordinates relative to lambda: AP +5.1 mm, ML 2.0 mm, and DV −1.5 mm, with an approximately 180-degree earbar-beak angle. Following the craniotomy at the target locations, coordinates were confirmed by electrophysiological recordings using 1 MΩ tungsten sharp electrodes. Subsequently, 200 nl of undiluted viral vector were injected into each hemisphere using glass pipettes attached to either a Nanoject III (Drummond Scientific), with a constant flow rate of 1 nl per second, or a custom built pressure injector with a similar flow rate. To avoid backflow of viral vector and untargeted transduction, injection pipettes were raised about 200 μm relative to the injection site after content delivery and let sit for 5 minutes prior to ejection. Cranial holes were covered with quickcast, and the incision site in the skin was closed with tissue glue.

Mice were deeply anesthetized with Ketamine (90 mg/kg body weight)/Xylazine (8 mg/kg body weight), with buprenorphine (0.1 mg/Kg body weight, i.p.) given pre-emptive 20 minutes prior to anesthesia. Once pedal-reflex was absent, mice were mounted into a stereotaxic frame (Kopf), and 300 nl of undiluted viral vector were injected unillaterally into the dorsal striatum (DS) or into area CA1 of the dorsal hippocampus at a rate of 50 nl/min employing a 33G needle (nanofil 33 G, WPI) in a 10 μl syringe (nanofil, WPI) and a microinjector pump (WPI; UMP3 UltraMicroPump) directly mounted to the stereotaxic manipulator. The needle was left in place for an additional 5 minutes after the completed injection to avoid backflow of the viral vector. After the injection needle was removed, the incision was sutured. Animals received analgesic treatment for 3 days after surgery. The following stereotaxic coordinates where used relative to bregma: AP +1.0 mm, ML +1.8 mm, and DV −3.0 mm for DS; and AP −2.0 mm, ML +1.5 mm, and DV −1.6 mm for CA1.

### Tissue preparation and immunohistochemistry

After isoflurane overdose, birds were transcardially perfused with PBS followed by 4% PFA (paraformaldehyde) in PBS (pH 7.4). Brains were extracted and post-fixed in 4% PFA overnight at 4 °C, stored in PBS, and hemisected. Hemispheres were sectioned into either 30 or 60 μm-thick sagittal sections using a freezing sliding microtome (Leica Microsystems).

9 to 10 days after viral vector injection, mice were anesthetized with a lethal dose of Pentobarbital (200 mg/kg, i.p.), and transcardially perfused with saline followed by 4% PFA. Brains were extracted and post-fixed in 4 % PFA for 24 hours. Sections were prepared with a Vibratome (Leica Microsystems) at 50 μm thickness.

Sections were either directly mounted onto glass slides with Vectashield antifade mounting medium (Vector Laboratories) and coverslipped for imaging of native fluorescence signal, or subjected to immunohistochemistry protocols, as described below.

Sections were washed with PBS, blocked in 10%normal goat serum/0.5% saponin/PBS for 90 minutes, and incubated overnight with primary antibodies in blocking solution at 4 °C. Sections were then washed in 0.5% saponin/PBS, and incubated with secondary antibodies in blocking solution for 3 hours at room temperature. Finally, sections were washed in 0.5% saponin/PBS, mounted onto glass slides with Vectashield antifade mounting medium (Vector Laboratories), and coverslipped.

The following primary antibodies were used: chicken anti-GFP (1:1000, Aves, GFP-1020), mouse anti-HuC/HuD (1:20, Invitrogen, A21271), and rabbit anti-TH (1:500, Millipore, AB152). The following secondary antibodies were used: goat anti-chicken conjugated to Alexa Fluor 488 (1:500, Abcam, ab150169), goat anti-mouse conjugated to Alexa Fluor 555 (1:500, Invitrogen, A21422), and goat anti-rabbit conjugated to Alexa Fluor 555 (1:500, Invitrogen, A21428).

### Image acquisition and analysis

Fluorescence images were acquired using a Leica SP8 confocal microscope (Leica Microsystems) using the 488 nm (eGFP/Alexa Fluor 488) and 561 nm (Alexa Fluor 555) laser line for excitation, and spectral windows of 493 nm - 550 nm and 566 nm - 650 nm respectively on hybrid detectors (HyD) for signal detection. Low magnification images were acquired using a 20x/0.75 NA multiimmersion objective (HC PL APO CS2), and higher magnification images with a 63x/1.3 NA glycerol-immersion objective (HCX PL APO CS2). Pixel resolution and z section distances were set considering the nyquist criteria as defined at https://svi.nl/NyquistCalculator to ensure optimal post processing results.

When imaging the entire HVC multiple field of views where imaged with a 10% overlap and merged after acquisition using Leica’s imaging software (LASX). Prior to analysis of image stacks recorded with the higher magnification objective images were deconvolved using Huygens professional (SVI). Spine reconstructions and density measurements were performed with Imaris 9.2.1 (Bitplane) using the semi-automatic filament tracing function. Soma counts and densities were performed using Imaris’ spot tool with user defined thresholds. Sections for analysis were selected in a way that they roughly represent the same anatomical location (i.e. similar medio-lateral positions), and we performed soma counts in standardized volumes of 300 x 300 x 30 μm^3^ to accommodate for slight variations in HVC size or section thickness.

### Calcium imaging

A custom-built, miniaturized fluorescence microscope was used to capture the temporal dynamics of calcium induced GFP signal from HVC projection neurons retrogradely transduced with our GCaMP6f-construct. The miniaturized microscope (named “finchscope”) was built in-lab according to the previously described manufacturing protocol^8^.

In order to image fluorescence signals, a cranial window was prepared in deeply anesthetized birds (by oral administration of a mixture of oxygen with 1-3% isoflurane) as follows. The skull and dura over HVC were removed, and a ~3 mm hole on the skull just above HVC was made using a dental drill and scalpel. A round cover glass (3-mm diameter, CS-3R-0, Warner Instruments) was placed on the brain surface and fixed to the skull with tissue adhesive (Gluture, World Precision Instruments). The edge of glass window was secured to the skull with dental cement (PalaXpress, Kulzer). We then recorded spontaneous calcium activity by placing the finchscope just above the glass window with a stereotaxic manipulator.

Movies were recorded with the finchscope at 30 FPS and (0.75 μm)^2^ pixel size via a video capture device (GV-USB2, I-O DATA) and a recording software (AMcap), and were processed with in-house developed MATLAB (Mathworks) scripts as follows. First, we temporally smoothed each pixel using a 5-frame moving average. Next, we removed wide field intensity fluctuations by dividing each frame by a low-pass-filtered version of itself. For further analysis, we re-expressed each pixel in units of relative changes in fluorescence, given by ΔF(t)/F0 = (F(t) - F0)/F0, where F0 is the mean pixel value calculated by averaging over the entire movie. To obtain activity traces and spatial filters corresponding to the imaged cells, we applied an established automated signal extraction method to the preprocessed movie^9^. Finally, we manually verified all extracted cells and discarded false positives.

